# Structural volume composition of internodes determines culm non-structural carbohydrates accumulation in rice

**DOI:** 10.64898/2026.01.19.700429

**Authors:** Yu Wakabayashi, Naohiro Aoki, Ryutaro Morita, Megumi Sudo, Yoichiro Kato

## Abstract

Non-structural carbohydrates (NSC) stored in the stem play a crucial role in supporting yield formation in rice. However, internode morphological determinants of NSC accumulation are unclear. This study aimed to clarify the relationship between internode morphology and NSC accumulation and to identify a robust morphological indicator for evaluating NSC accumulation capacity. Two years of field experiments were conducted using multiple cultivars. The NSC content was quantified for individual internodes and at the whole-plant culm level, and its relationships with internode morphological traits were analyzed. Since the upper internodes (UIN; first and second internodes) and lower internodes (LIN; third and subsequent internodes) exhibited contrasting roles in NSC accumulation, a novel index was introduced, the volume composition ratio (VCR) of UIN/LIN, which represents their relative volumetric contributions within a culm. The VCR of UIN/LIN showed the strongest correlation with culm NSC and high reproducibility across years, outperforming simple morphological traits. Manipulation of internode development using plant growth regulators demonstrated that altering VCR effectively modified culm NSC accumulation. Accordingly, the VCR of UIN/LIN serves as a robust morphological indicator of culm NSC accumulation capacity, providing a practical framework for improving NSC accumulation to achieve high and stable yield performance in rice.

**Highlight:** This novel internode structural index robustly predicts the culm non-structural carbohydrate accumulation capacity, providing a practical morphological indicator for improving yield stability in rice.

## Introduction

Rice (*Oryza sativa* L.) is a staple food for nearly half of the world’s population and accounts for approximately 17% of the global caloric intake (FAO 2023; IRRI, 2016). Therefore, increasing rice production is of great significance for global food security to feed the rapidly increasing world population (Khush, 2005; Powell *et al*., 2012).

To date, numerous high-yielding rice cultivars with extra-large sink capacities, characterized by a high harvest index and increased spikelet number per panicle, have been developed, such as the IRRI’s New Plant Type and Chinese hybrid rice (Khush, 2013;Peng *et al*., 1999). In Japan, many high-yield cultivars with extra-large sink capacities, including Takanari, Momiroman, and Hokuriku193, have been developed since the early 1980s (Taylaran *et al*., 2009; Yoshinaga *et al*., 2013). These breeding efforts have focused on improving sink-related traits, such as spikelet number and grain weight. Although studies using these cultivars have contributed to the identification of many genes involved in sink capacity (Chen *et al*., 2021; Lu *et al*., 2022), sink expansion alone does not necessarily lead to improved grain yield (Fukushima *et al*., 2017; Nakano *et al*., 2017; Ohsumi *et al*., 2011) and enhancement of carbohydrate supply ability (source capacity) must be pursued in parallel (Ueda *et al*., 2025).

Carbohydrates used for grain development originate from two major sources: carbohydrates stored in the stem prior to heading and photosynthates newly-produced during the grain-filling period. Under normal growing conditions, approximately 25% of the final grain mass is derived from the former, with the remainder supplied by the latter (Cock and Yoshida, 1972). The stem serves as a major temporary storage organ that accumulates soluble sugars and starch, which can be rapidly remobilized to the grains after heading (Perez *et al*., 1971). These temporarily-stored carbohydrates are referred to as non-structural carbohydrates (NSC) to distinguish them from structural carbohydrates such as cellulose and lignin, which constitute cell wall components.

Within a rice panicle, inferior spikelets, which are located on the lower branches and flower later, are characterized by slow and incomplete grain filling (Mohapatra *et al*., 1993;Zhao *et al*., 2020). Enhancing the grain-filling ability of inferior spikelets is particularly important for high-yielding varieties, in which the large sink size often exceeds the available source capacity (You *et al*., 2016). NSC reserves play a key role in determining the grain-filling performance of inferior spikelets in such varieties, and increasing NSC availability largely contributes to higher grain yield (Chen *et al*., 2019; Fu *et al*., 2011;Lin *et al*., 2024). In addition, NSC reserves are known to mitigate yield losses when exposed to unfavorable conditions during grain filling (Slewinski, 2012). Under stresses such as low solar radiation (Nagata *et al*., 2001; Okamura *et al*., 2013;Okawa *et al*., 2003), low nitrogen availability (Ju *et al*., 2015;Li *et al*., 2018; Pan *et al*., 2011), drought (Yang *et al*., 2001; Yang and Zhang, 2010), and high temperature (Morita and Nakano, 2011), the relative contribution of stem NSC to grain filling increases.

Although NSC accumulation at the pre-heading stage is strongly influenced by environmental factors, such as soil nitrogen status (Ju *et al*., 2015;Li *et al*., 2018; Pan *et al*., 2011) and solar radiation (Nagata *et al*., 2001), substantial genotypic variation in NSC accumulation has been reported (Samonte *et al*., 2001; Lubis *et al*., 2003; Arai-Sanoh *et al*., 2013; Yoshinaga *et al*., 2013; Tu *et al*., 2025). Some varietal differences have been explained by biomass production capacity (Takai et al., 2006), growth duration (Nagata *et al*., 2002), and physiological factors (Hirano *et al*., 2005; Wang *et al*., 2017; Okamura *et al*., 2018). However, no study has systematically evaluated the relationship between culm morphological traits and the NSC accumulation capacity. Previously, we analyzed NSC dynamics across nodal positions at the pre- and post-heading stages in two large-panicle cultivars differing in NSC accumulation at heading (Wakabayashi *et al*., 2022). We found clear genotypic differences in NSC accumulation, specifically in the lower internodes of the culm, rather than in the upper internodes or leaf sheaths. Our study suggests that these genotypic differences are associated with internode morphological traits, such as length and diameter, although the specific causal factors and their generality across cultivars were unclear.

Therefore, in the current study, a multi-cultivar comparison was conducted to identify the morphological factors associated with NSC accumulation in the culm. Furthermore, because manipulating culm morphology using plant growth regulators provides an effective approach in assessing causal relationships, the relevant morphological traits were experimentally modified and how these changes affected NSC accumulation was investigated.

## Materials and methods

### Experimental design

Two field experiments and one pot experiment were conducted. The 2022 field experiment was designed to examine the relationship between NSC accumulation and morphological traits of each internode in the main culm. The 2024 field experiment aimed to relate plant-level culm NSC accumulation to the morphological traits of each internode in the main culm under more uniform climatic conditions across cultivars during the period from panicle formation to heading when NSC accumulates in the culm. A pot experiment was conducted to evaluate how altering the internode composition within a culm by applying plant growth regulators (gibberellin and uniconazole) affected NSC accumulation.

### Field experiments

Field experiments were conducted in 2022 and 2024 in the paddy field at the Institute for Sustainable Agro-ecosystem Services (ISAS), The University of Tokyo (35°44′N, 139°32′E, altitude: 58 m), Tokyo, Japan. Ten high-yielding cultivars with different genome types were used (Supplementary Table S1). Seeds were sown in seedling nursery boxes, and seedlings (approximately 25 d old) were transplanted on 31 May 2022. In 2024, the sowing and transplanting dates were adjusted for each cultivar based on the heading dates in 2022 to synchronize the heading period. In both years, the planting density was 22.2 hills m^-2^ (300-mm row spacing, 150-mm hill spacing). Three seedlings per hill were transplanted in two rows (15 plants per row, 2.1 m) for each cultivar. N (6 g m^-2^ as urea), P (8 g m^-2^ as single superphosphate), and K (9 g m^-2^ as potassium chloride) were applied as basal fertilizer before transplanting. N (4 g m^-2^ as ammonium sulfate) and P (3 g m^-2^ as single superphosphate) were applied as top dressings at the tillering stage on 12 July 2022, and 20 July 2024. Weeds, insects, and diseases were prevented by administering herbicides and pesticides as needed. The plots were arranged in a randomized block design with four replicates. The cultivar arrangement in the plot was fully randomized in 2022, whereas two cultivars with similar transplant dates were randomly placed next to each other in 2024.

In 2022, four main stems were sampled from different plants at 5 d after heading (DAH), when the first internode had completed elongation. One stem whose culm length was closest to the mean of the four sampled stems was selected as a representative. The selected stems were divided into individual internodes and leaf sheaths, and the lengths of each internode and leaf sheath were measured. An approximately 1 mm-thick cross-section was cut at the midpoint of each internode using a razor blade, stained with safranin, and scanned using a GT-X980 scanner (Epson, Nagano, Japan) for cross-sectional trait analysis. Internodes at each position (including the decolorized section prepared for cross-sectional imaging), leaf sheaths, leaf blades, and panicle were dried at 80 °C for at least 72 h and weighed. Each internode and leaf sheath was ground to a powder using a ShakeMaster Auto (Biomedical Science, Tokyo, Japan) for soluble sugar and starch measurements.

In 2024, three whole plants per plot were sampled at full heading, when 80–90% of the panicles had fully emerged, and dried at 80 °C for at least 72 h. One plant whose aboveground dry weight was closest to the mean was selected as a representative. The plant was divided into panicles, leaf blades, culms, and leaf sheaths, and each organ was weighed. Culms and leaf sheaths were ground to powder using a Wiley laboratory mill (1029-C, Yoshida Seisakusho, Tokyo, Japan) and a Hi-speed vibration sample mill (TI-200, Cosmic Mechanical Technology, Fukushima, Japan) for soluble sugar and starch measurements. Three main stems per plot were sampled and one stem with a culm length closest to the mean was selected. The selected stem was divided into individual internodes, and the internode length and cross-sectional traits were measured as in 2022. To assess the NSC-buffering effect under source-limited conditions during the grain-filling period, all flags and second leaves were manually removed from the four plants per plot at full heading. At maturity, panicles from four plants with or without defoliation treatment were harvested and threshed, and their dry weights were determined after drying at 80 °C for at least 72 h. Total spikelet numbers per plant were counted in plants without defoliation treatment.

### Pot experiment

Pot experiment was conducted in a greenhouse at ISAS using the cultivar ‘Hokuriku 193’, one of the highest-yielding cultivars in Japan (Okamura et al., 2022). Seeds were sown in a seedling nursery box on 5 May, and ten seedlings were transplanted into 1/5000-a Wagner pots in a circular arrangement on 1 June 2024. All emerging tillers were removed throughout the growth period so that only the main stem developed. Plant growth regulator treatments were applied at 25 d before heading (DBH) and 3 DBH, corresponding to the periods of lower and upper internode elongation, respectively (Wakabayashi *et al*., 2022).

Seven treatment groups were established based on a combination of application timing and chemical components (Supplementary Fig. S1A). A 100-μL aliquot of either 70 μM gibberellin solution (GAL dissolved in 0.6% [v/v] ethanol) or 100 μM uniconazole solution (uniconazole-P dissolved in 0.6% [v/v] ethanol) was applied to each plant for three consecutive days. The control plants were treated with a mock solution containing 0.6% (v/v) ethanol. The solutions were injected into the gap between the leaf sheaths using a micropipette (Supplementary Fig. S1B). To affect the elongation of the fourth internode (IN4), the leaf sheath attached to the fourth leaf blade below the uppermost expanding leaf was targeted on 28–30 July. To affect the elongation of the second internode (IN2), the second leaf sheath was treated on 23–25 August.

At 0 DAH, when the first and second internodes (IN1 and IN2) were elongating, internodes IN2 and IN4 and their corresponding leaf sheaths (LS2 and LS4) were sampled from five plants and immediately frozen in liquid NL. These samples were ground into a powder using a ShakeMaster Auto for ADP-glucose pyrophosphatase (AGP) activity assays. At 5 DAH, five plants were sampled and divided into panicles, leaf blades, culms, and leaf sheaths. The culm and leaf sheaths were further divided into individual nodal segments. Internode length, cross-sectional traits, and leaf sheath length were measured as in the field experiments. Each organ was dried at 80 °C for at least 72 h and weighed. Each internode and leaf sheath was ground to a powder for soluble sugar and starch measurements.

### Analysis of images obtained using a scanner

The cross-sectional traits (diameter, wall thickness, and cross-sectional area) of each internode were analyzed from the scanned images (Supplementary Fig. S2). The areas of the outer and inner circles were measured using ImageJ software (ver. 1.54g). Each cross-sectional trait was calculated using the following equation:

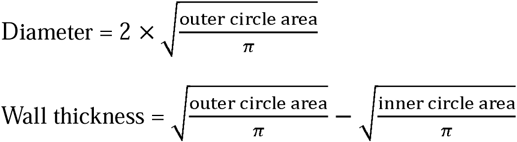

Cross-sectional area = outer circle area – inner circle area

Volume = internode length × cross-sectional area

### Measurement of starch and soluble sugars

Soluble sugar (sucrose, glucose, and fructose) and starch contents were measured according to the method of Okamura *et al*. (2013) using glucoamylase (Toyobo Co., Osaka, Japan), an E-kit (E1247A; J.K. International, Tokyo, Japan), and a microplate reader (Synergy H1; Agilent Technologies, California, USA).

### Enzyme activity assay

The activity of ADP–glucose pyrophosphorylase (AGP; EC 2.7.7.27) was analyzed because it catalyzes the first committed step of starch biosynthesis and is commonly used as a proxy for sink strength in rice stems (Okamura *et al*., 2013, 2018; Jiang *et al*., 2022). Approximately 50 mg of the ground sample was suspended in 10 volumes of extraction buffer (50 mM HEPES–KOH (pH 7.4), 5 mM MgClL, 2 mM EDTA, 12.5% [*v*/*v*] glycerol, 5 mM DTT, 5% [*w*/*v*] polyvinylpolypyrrolidone, and one tablet of protease inhibitor cocktail [cOmplete Mini, EDTA-free] per 10 mL). After centrifugation at 12,000 × *g* for 5 min at 4 °C, the supernatant was used as the crude enzyme extract for the AGP assays.

The AGP activity assay was conducted in reaction mixtures consisting of 20 µL crude enzyme sample and 174 µL reaction solution containing 50 mM HEPES–KOH (pH 7.4), 3 mM 3-phosphoglycerate, 4 mM pyrophosphate, 5 mM MgCl_2_, 4 mM DTT, 0.8 mM NADP^+^, 0.5 U phosphoglucomutase, and 1 U glucose-6-phosphate dehydrogenase. The reaction was initiated by the addition of 6 µL of ADP-glucose (final concentration, 3 mM), and the increase in absorbance at 340 nm was monitored at 30 °C for 10 min using a microplate reader. The AGP activity was calculated as the difference between the reaction rate after the addition of ADP–glucose and the pre-addition baseline rate.

### Statistical analysis

All statistical analyses were performed using R software (v.4.0.3). Cultivar effects on NSC-related and morphological traits were evaluated using a one-way analysis of variance (ANOVA), followed by Tukey’s honestly significant difference (HSD) test for multiple comparisons among cultivars. In the growth-regulator experiment, MM (mock-treated plants receiving solvent only) was used as the control and the differences between MM and each treatment were assessed using Dunnett’s test. The relationships between NSC-related and morphological traits were examined using Pearson’s correlation coefficients. All the hypothesis-based statistical tests were two-sided. Hierarchical clustering was performed using Ward’s method on a correlation matrix. In all analyses, *n* indicates the number of biological replicates (for individual plants).

## Results

### NSC accumulation traits

In the 2022 field experiment, NSC accumulation traits were evaluated in the main culm to characterize cultivar differences in each internode. Although all cultivars are generally classified as heavy-panicle types, they differed in panicle number and heading period (Supplementary Fig. S3; Supplementary Table S1). Therefore, in the 2024 experiment, NSC accumulation in culms was evaluated at the whole-plant level, with sowing dates adjusted to minimize the variation in heading time (Supplementary Table S1). Compared with the 2022 experiment, cultivar differences in air temperature and solar radiation during the pre-heading period were smaller in the 2024 experiment (Supplementary Fig. S4). Culm NSC content showed no correlation with air temperature or solar radiation during the pre-heading stage (Supplementary Fig. S5), indicating that genetic factors were the primary drivers of NSC variation.

Significant cultivar differences were observed in the main culm NSC content in 2022 (Fig. 1A), and similar differences were detected at the whole-plant level in 2024 (Fig. 1B, C), indicating that NSC accumulation in the main culm reflected that at the whole-plant level. NSC content also differed significantly among the internodes (IN1-IN5+), with greater variation in IN4 and IN5+ than in the upper internodes (Supplementary Table S2). To further characterize NSC accumulation, the NSC content was partitioned into two components: the residual components (RC), calculated by subtracting NSC from organ dry weight, and NSC/RC, obtained by dividing NSC by RC. RC represents the amount of dry matter invested in an organ that is not utilized for NSC accumulation, whereas NSC/RC represents the amount of NSC per unit of invested RC, indicating the efficiency of NSC accumulation. Both RC and NSC/RC also showed significant cultivar differences at every internode as well as in the main and whole-plant culms (Supplementary Table S2).

**Fig. 1.**
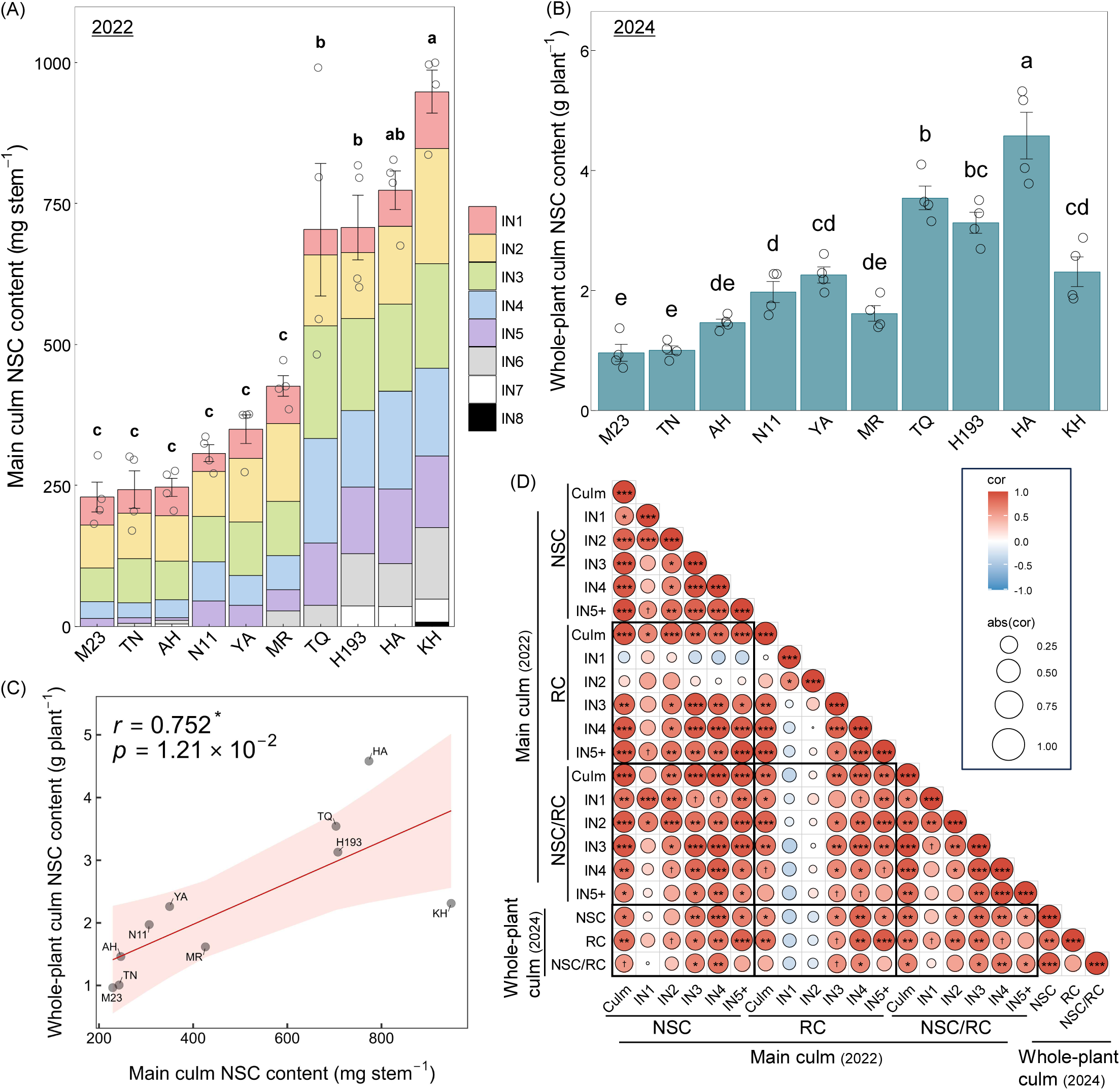
Culm NSC accumulation and its relationships with morphological and NSC-related traits for each cultivar. (A) NSC content of the main culm at heading in 2022. Bars represent stacked NSC contents of individual internodes (IN1–IN8). Bars show means ± SE (*n* = 4). Different letters above bars indicate significant differences among the cultivars (Tukey’s HSD, *P* < 0.05). (B) NSC content at the whole-plant culm level in 2024. Bars show means ± SE (*n* = 4). Different letters above bars indicate significant differences among the cultivars (Tukey’s HSD, *P* < 0.05). (C) Relationship between the main culm NSC content (2022) and the whole-plant culm NSC content (2024). Regression lines are shown with 95% confidence intervals (shaded). Correlation coefficients (*r*) and *P*-values are from Pearson’s tests (*n* = 10). (D) Correlation matrix among NSC-related traits (NSC, RC, NSC/RC) of individual internodes (IN1–IN5+) in the main culm or whole-plant culms. IN5+ indicates internodes at and below IN5. Circle size and color indicate correlation strength and direction, respectively (Pearson’s *r*). Significance levels: † *P* < 0.10, * *P* < 0.05, ** *P* < 0.01, *** *P* < 0.001 (*n* = 10).

Correlation analyses indicated that RC in IN3–IN5+ was positively associated with NSC in IN2–IN5+, whereas RC in IN1 and IN2 showed no such relationship with any of the internodes (Fig. 1D). NSC/RC was strongly correlated with NSC in the same or adjacent internodes within a culm. At the whole-plant level, NSC was positively correlated with both RC and NSC/RC.

Leaf sheath NSC also differed among cultivars, and showed strong positive correlations with culm NSC except in two cultivars (MR and HA) (Supplementary Fig. S6). The regression slopes indicated that NSC accumulation in leaf sheaths reached one-third to one-half of the culm NSC content, reflecting parallel variation between the two organs while differing in the extent of NSC accumulation. In addition, culm NSC, but not leaf-sheath NSC, was negatively correlated with the reduction ratio of grain dry weight under defoliation, indicating that culm NSC contributed more strongly to mitigating yield loss than leaf-sheath NSC (Supplementary Fig. S7).

### Morphological traits

To evaluate cultivar differences in the morphological traits of each internode, internode volume, length, cross-sectional area, diameter, and wall thickness were quantified. Significant cultivar differences were observed in internode volume across all internodes in both years, with greater variation observed in the lower internodes (Fig. 2B; Supplementary Table S3, S4). Most other morphological traits showed significant cultivar differences, except for the cross-sectional areas of IN1 and IN5+ and the wall thickness of IN2 in 2022. Correlation analyses indicated that internode length was positively associated with volume in all internodes in 2022 and in IN3–IN5+ in 2024 (Fig. 2C). The cross-sectional area was positively correlated with the volume in IN2 in both years. The diameter exhibited a consistently stronger relationship with the cross-sectional area than did the wall thickness. In particular, the diameters of IN2–IN4 were significantly correlated with the cross-sectional area in both years.

**Fig. 2.**
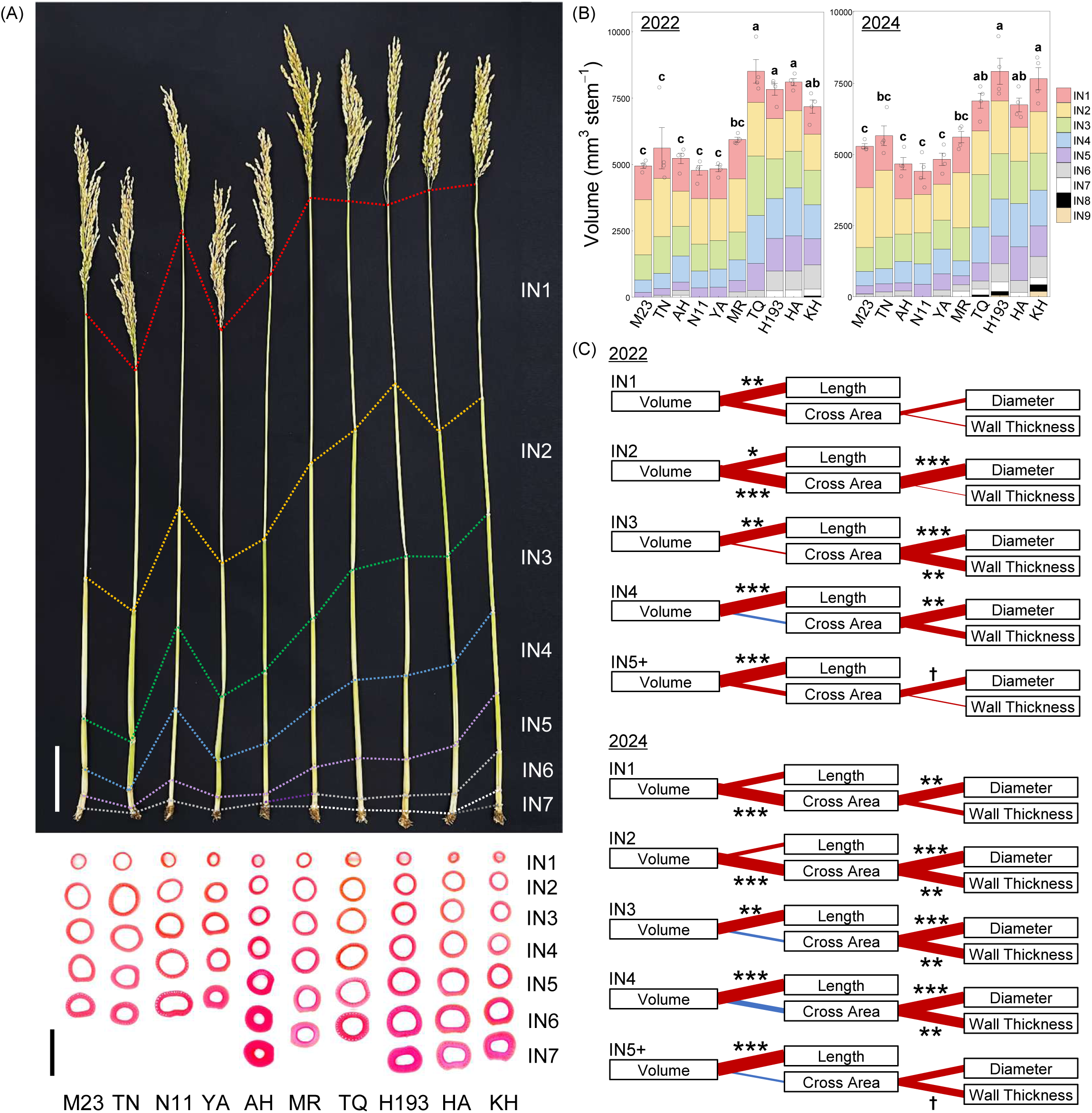
Morphological traits of individual internodes for each cultivar. (A) Main culm at heading for each cultivar. Each internode position is indicated by a colored dashed line. The bottom panel shows cross-sectional images of each internode, aligned by cultivar and nodal position, indicating genotypic variation in internode diameter and wall thickness. Scale bars indicate 100 mm for the whole culm (top) and 10 mm for the transverse cross-sections of internodes (bottom). (B) Internode volume for each cultivar in 2022 (left) and 2024 (right). Bars represent stacked volumes of the individual internodes. Bars show means ± SE (*n* = 4). Different letters above bars indicate significant differences among the cultivars (Tukey’s HSD, *P* < 0.05). (C) Relationships between internode volume and its component traits (internode length, cross-sectional area, diameter, and wall thickness) for each internode position (IN1–IN5+) in 2022 and 2024. Red or blue lines indicate positive or negative correlations, respectively. Line thickness corresponds to correlation strength. Significance levels: † *P* < 0.10, * *P* < 0.05, ** *P* < 0.01, *** *P* < 0.001 (*n* = 10).

### Morphological factors associated with NSC accumulation

To clarify the relationship between morphological factors and NSC accumulation, the correlations between internode volume and NSC-related traits at each internode were analyzed (Fig. 3A). The volumes of IN1 and IN2 had no significant correlation with NSC, RC, or NSC/RC in most internodes, indicating a minimal contribution of the upper internodes to culm NSC accumulation. In contrast, the volumes of IN3–IN5+ were positively correlated with NSC and RC in most of IN3–IN5+ as well as in the culm, and with NSC/RC across multiple internodes. Therefore, IN3–IN5+ play a major role in determining culm NSC accumulation. Cluster analysis indicated that the RC and volumes of IN1 and IN2 formed a distinct cluster separate from other traits, including culm NSC (Fig. 3B). Based on these results, IN1 and IN2 were defined as upper internodes (UIN) and IN3–IN5+ as lower internodes (LIN). Both UIN and LIN volumes showed significant cultivar differences, with the variation being larger in LIN than in UIN in both years (Fig. 3C). Cultivar differences were also observed in the volume composition ratio (VCR) of UIN and LIN within the culm.

**Fig. 3.**
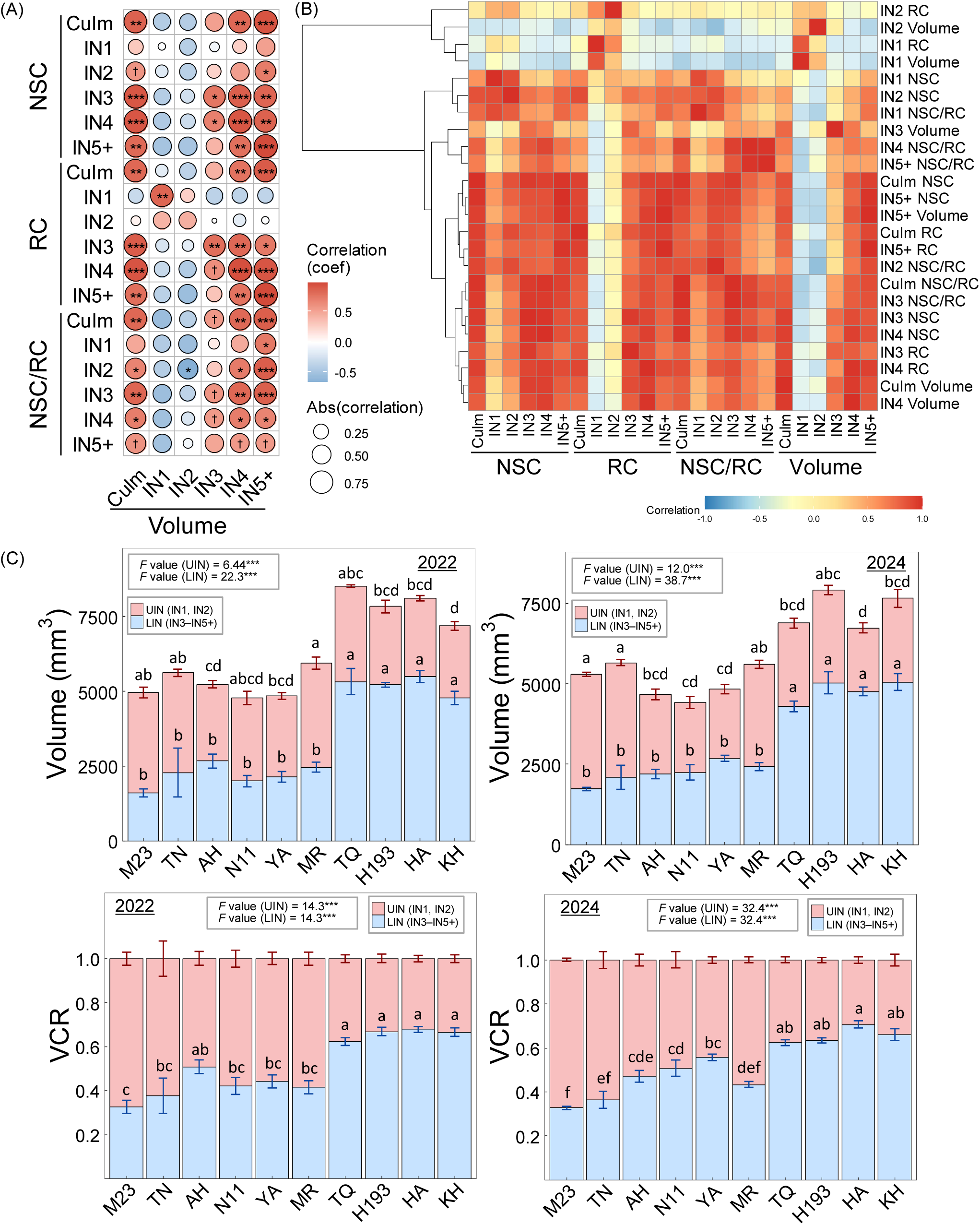
Relationships among NSC-related traits and internode morphological characteristics across cultivars. (A) Correlation matrix between the internode volume and NSC-related traits (NSC, RC, NSC/RC) for each internode (IN1–IN5+). (B) Heatmap of pairwise correlations among the NSC, RC, NSC/RC, and internode volumes in 2022. Hierarchical clustering (Ward’s method) highlights trait groups showing similar correlation patterns across the cultivars. (C) Cultivar differences in the internode volume (top) and VCR (bottom) for UIN (IN1–IN2) and LIN (IN3–IN5+) in 2022 (left) and 2024 (right). Bars show means ± SE (*n* = 4). Different letters above bars indicate significant differences among the cultivars (Tukey’s HSD, *P* < 0.05). *F* values and significance levels from the ANOVA are shown in each panel. Significance levels: *** *P* < 0.001. The VCRs of UIN and LIN are complementary components of a proportional index that sums up to one within a culm. Therefore, both components exhibit identical *F* values and share the same groupings in multiple-comparison tests.

To further identify the morphological factors most strongly associated with NSC accumulation, the relationships between NSC content and morphological traits were examined with their year-to-year reproducibility (Fig. 4A). In three relationships— between main-culm morphological traits and either main culm NSC or whole-plant NSC, and the year-to-year reproducibility of each trait— the VCR in IN1, IN2, IN4, IN5, UIN and LIN, the volumes in IN4 and IN5+, and the length in IN3 and IN4 showed significant correlations. In particular, the VCR in UIN and LIN (complementary traits that sum up to one within a culm) showed a relatively strong correlation in each relationship (Fig. 4B; Supplementary Fig. S8), indicating that these traits were less susceptible to environmental factors and were the most suitable indicators for evaluating cultivar differences in terms of NSC accumulation.

**Fig. 4.**
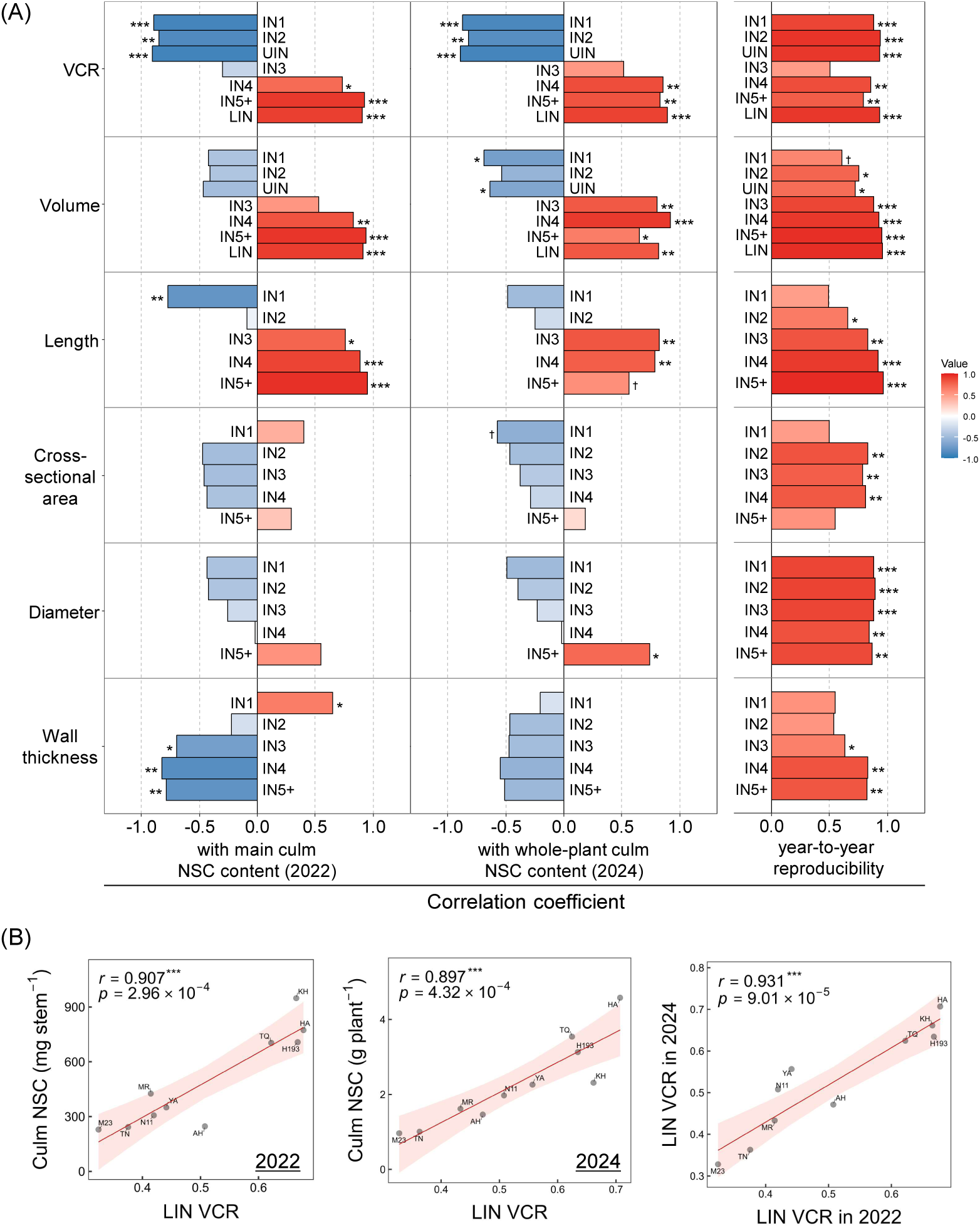
Relationships between internode morphological traits and culm NSC accumulation. (A) Correlation coefficients between internode morphological traits (VCR, volume, length, cross-sectional area, diameter, and wall thickness) and main culm NSC content in 2022 (left), whole-plant culm NSC content in 2024 (middle), and year-to-year reproducibility of each trait (right). Bars represent Pearson’s correlation coefficients for each internode (IN1–IN5+), including UIN (IN1–IN2) and LIN (IN3–IN5+). Red and blue bars indicate positive and negative correlations, respectively. Significance levels: † *P* < 0.10, * *P* < 0.05, ** *P* < 0.01, *** *P* < 0.001 (*n* = 10). (B) Relationships between LIN VCR and main culm NSC content in 2022 (left), between LIN VCR and whole-plant culm NSC content in 2024 (middle) and between LIN VCR in 2022 and 2024 (right), showing year-to-year stability. Regression lines are shown with 95% confidence intervals (shaded). Correlation coefficients (*r*) and *P*-values are from Pearson’s tests (*n* = 10).

### Effect of VCR-altering treatment on NSC accumulation

Based on the results of the field experiments using multiple cultivars, it was hypothesized that the VCR of UIN/LIN within a culm was a key factor in determining NSC accumulation. To test this, gibberellin (GA) and uniconazole (UZ) treatments were applied to modify the VCR in Hokuriku 193 (Fig. 5A). GA and UZ treatments targeting IN4 significantly increased or decreased LIN volumes, respectively, except in UG (Fig. 5B). Treatments targeting IN2 increased or decreased the UIN volumes, except in GU. Consequently, the VCR of UIN or LIN was altered in each treatment group, with most changes explained by alterations in the internode length rather than in the cross-sectional area (Fig. 5C).

**Fig. 5.**
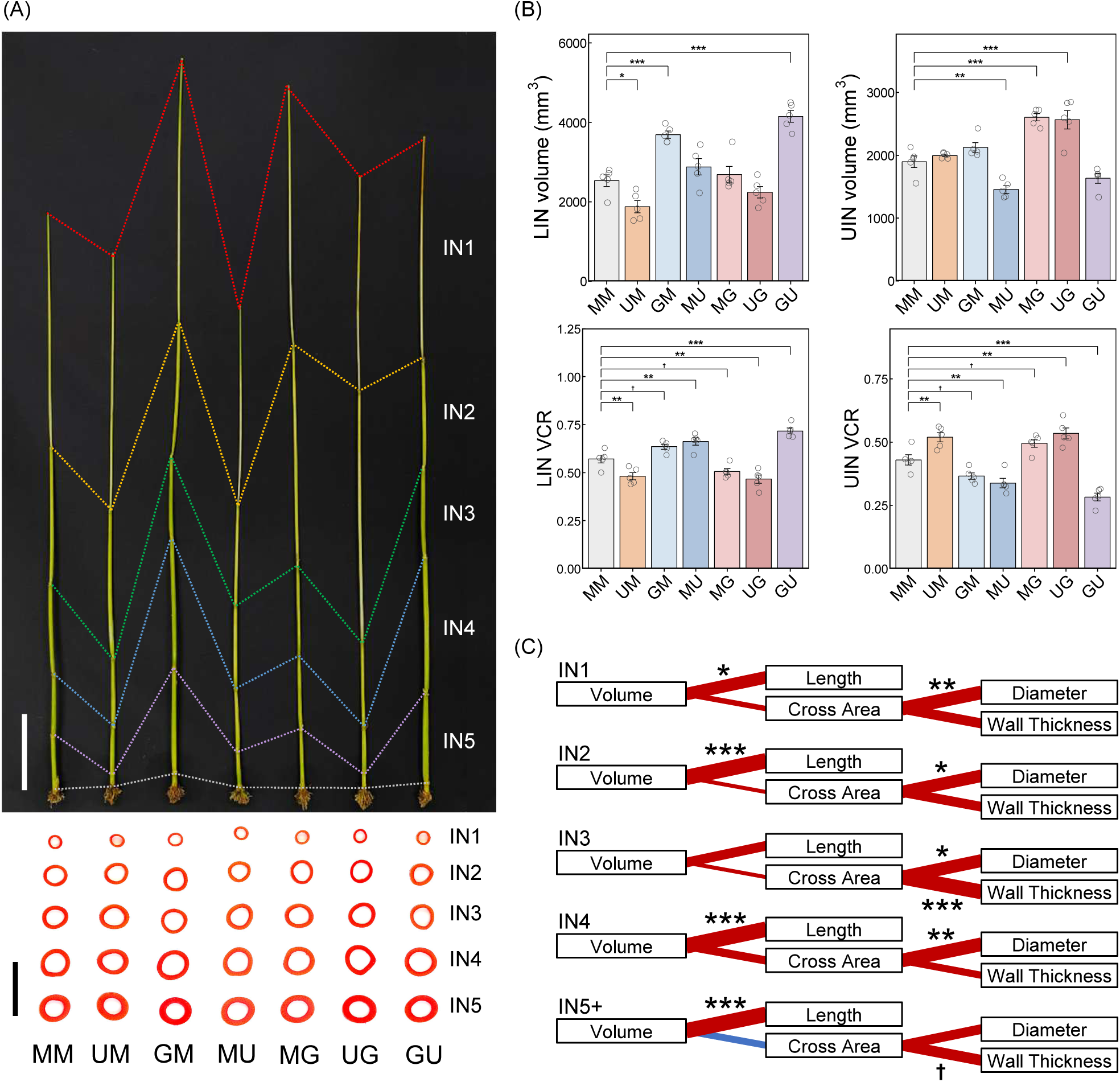
Effects of plant growth regulator treatments on internode morphology and VCR of the main culm. (A) Main culm for each treatment group (MM, UM, GM, MU, MG, UG, GU). Each internode position is indicated by a colored dashed line (IN1–IN5). The bottom panel shows cross-sectional images of each internode, aligned by treatment and nodal position, indicating treatment variation in internode diameter and wall thickness. Scale bars indicate 100 mm for the whole culm (top) and 10 mm for the transverse cross-sections of internodes (bottom). (B) Effects of gibberellin (GA) and uniconazole (UZ) on UIN (IN1–IN2) and LIN (IN3–IN5+) morphology. Volume (top) and volume composition ratio (VCR) (bottom) of UIN and LIN are shown. Bars show means ± SE (*n* = 5). Horizontal brackets indicate statistically significant differences among treatments (Dunnett vs. MM, * *P* < 0.05, ** *P* < 0.01, *** *P* < 0.001). (C) Correlations between internode volume and morphological traits (length, cross-sectional area, diameter, and wall thickness) for IN1–IN5+. Red or blue lines indicate positive or negative correlations, respectively. Line thickness corresponds to correlation strength. Significance levels: † *P* < 0.10, * *P* < 0.05, ** *P* < 0.01, *** *P* < 0.001 (*n* = 7).

The culm NSC content increased significantly in GU and decreased in UM and UG relative to MM (Fig. 6A). Although not statistically significant, NSC tended to be higher in GM and MU and lower in MG. The UIN volume was not correlated with culm NSC, whereas the LIN volume was positively correlated (Fig. 6B). The VCR of UIN or LIN within a culm showed strong correlations with culm NSC, which was consistent with the field experiments (Fig. 4B, 6B). NSC-related traits were examined in the UIN and LIN to further clarify the basis of these changes (Fig. 6C). The NSC of UIN was significantly higher in GM, accompanied by increases in both RC and NSC/RC, despite no changes in the UIN volume. An increase in NSC/RC was also observed in GU, indicating that GA applied to IN4 influenced NSC accumulation in the upper internodes. No decline in UIN NSC was observed in MG or UG, although both treatments showed increases in RC and reductions in NSC/RC. In LIN, NSC decreased in UM and UG due to reductions in RC. Although not statistically significant, MG and UG showed a slight decline in NSC/RC, suggesting that enhanced elongation of the upper internodes may have limited NSC accumulation in the lower internodes. In contrast, GU exhibited an increase in the lower-internode NSC content due to a large increase in RC, despite the reduced NSC/RC. GM showed no increase in NSC because the increase in RC was offset by a substantial decrease in NSC/RC. Although the effects were not statistically significant, MU showed a slight increase in both RC and NSC/RC, which may account for the differences in NSC levels between GM and GU. Morphological changes in the internodes affected NSC accumulation not only in the culm, but also in the leaf sheaths (Supplementary Fig. S9). Leaf sheath NSC decreased significantly in MG and UG, mainly due to a reduction in NSC in the lower leaf sheaths (LS3–LS6) rather than the upper leaf sheaths (LS1 and LS2).

**Fig. 6.**
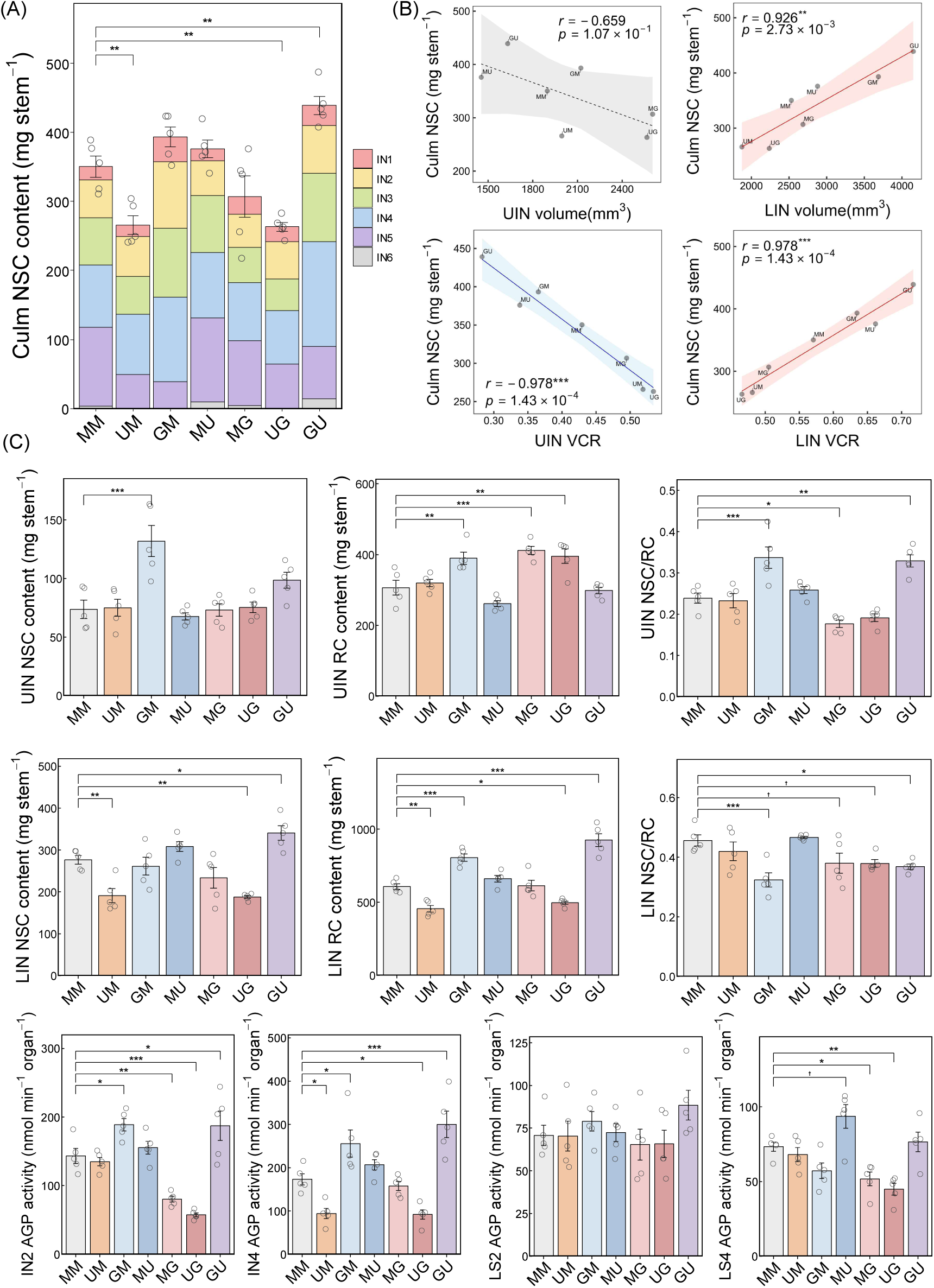
Effects of VCR-modifying treatments on culm NSC accumulation and NSC-related traits in upper and lower internodes. (A) Culm NSC content at 5 d after heading for each treatment (MM, UM, GM, MU, MG, UG, GU). Bars represent stacked NSC contents of individual internodes (IN1–IN5+). Bars show means ± SE (*n* = 5). Horizontal brackets indicate significant differences among treatments (Dunnett vs. MM, ** *P* < 0.01). (B) Relationships between culm NSC content and the volume of UIN (IN1–IN2) (top left) and LIN (IN3–IN5+) (top right), as well as the VCR of UIN (bottom left) and LIN (bottom right). Regression lines are shown with 95% confidence intervals (shaded). Correlation coefficients (*r*) and *P*-values are from Pearson’s tests (*n* = 7). (C) NSC-related traits in UIN and LIN for each treatment. NSC content, RC content, and NSC/RC in UIN (Top) and LIN (Bottom) are shown. Bars show means ± SE (*n* = 5). Horizontal brackets indicate statistically significant differences among the treatments (Dunnett vs. MM, †*P* < 0.10, * *P* < 0.05, ** *P* < 0.01, *** *P* < 0.001).

The AGP activity per organ was determined as an indicator of sink strength in IN2, IN4, LS2, and LS4 (Fig. 6D). AGP activity in IN2 increased significantly in GM and GU and decreased in MG and UG, largely consistent with the changes in NSC/RC in these treatments. In IN4, AGP activity increased in GM and GU, but decreased in UM and UG. These results indicate that elongation of the upper internodes reduced the sink strength within the same internode, whereas elongation of the lower internodes exerted an opposing effect. No significant differences were observed in LS2. In contrast, AGP activity at LS4 decreased significantly in MG and UG and slightly increased in MU, indicating that lower leaf sheaths were more sensitive to changes in upper internode morphology than upper leaf sheaths.

## Discussion

### Contrasting roles of upper and lower internodes in culm NSC storage

The relationship between NSC accumulation and various morphological traits in the culm were examined and clearly demonstrated that the volume composition ratio (VCR) of the internodes was a key determinant of NSC accumulation capacity. The contribution of internode volume to NSC accumulation differed markedly between the upper internodes (UIN: IN1 and IN2) and lower internodes (LIN: IN3–IN5+). The volume of UIN showed no or even a negative association with NSC accumulation (Fig. 3A). In particular, IN2 volume was negatively correlated with NSC accumulation efficiency (NSC/RC) in IN2, and this negative relationship was supported by the experiment with the plant regulator treatment (Fig. 6C, D). During rapid internode elongation, a substantial proportion of assimilated carbon is invested in structural growth, as evidenced by the strong accumulation of cell wall components in the elongating internodes (Lin *et al*., 2017). Therefore, assimilate investment for internode elongation and structural growth is likely to compete with NSC accumulation (Fujita and Yoshida, 1984; Arai-Sanoh *et al*., 2011; Botha *et al*., 2023), and reduce the efficiency of these internodes as storage organs.

In contrast, the volume of LIN was a strong determinant of culm NSC accumulation (Fig. 3A). Culms with larger LIN volumes exhibited greater NSC storage capacities, which is consistent with previous findings in wheat (Ehdaie *et al*., 2006a). These larger LIN volumes were primarily attributable to increased internode length rather than cross-sectional traits (Fig. 2C), as in sweet sorghum (Bihmidine *et al*., 2015). Our previous study indicated that the LIN function as major NSC storage organs during the pre-heading stage because they begin elongating earlier and act as sinks sooner than the UIN (Wakabayashi *et al*., 2022). Therefore, this temporal advantage likely enables the larger LIN to accumulate NSC more efficiently (Botha and Marquardt, 2024), even though the larger LIN require greater carbon investment for their elongation and structural formation, as observed for the UIN. The positional differentiation of internodes in carbohydrate storage has also been reported in other crops, including wheat (Ehdaie *et al*., 2006b; Li *et al*., 2013; Chen *et al*., 2014; Joudi *et al*., 2024), sweet sorghum (Hoffmann-Thoma *et al*., 1996; Gutjahr *et al*., 2013; Bihmidine *et al*., 2015; Shukla *et al*., 2017; Kanbar *et al*., 2021), and sugarcane (Wu and Birch, 2007; Vasantha *et al*., 2022; Botha *et al*., 2023; Botha and Marquardt, 2024), in which mature lower internodes preferentially accumulate carbohydrates. In grasses, internode elongation is generally initiated in the lower internodes and proceeds acropetally. Accordingly, the internode volume composition reflects not only the spatial architecture but also the temporal hierarchy of sink establishment within grass culms.

Despite the functional differentiation between UIN and LIN, the LIN exerted positive effects not only on their own NSC storage but also on NSC accumulation efficiency in the UIN, as indicated by the positive associations between the NSC/RC of the UIN and the residual components (RC) or volume of the LIN (Fig. 1D, 3A). This observation was supported by the plant growth regulator experiment, in which the elongation of LIN enhanced sink strength in UIN, thereby increasing their NSC accumulation (Fig. 6C, D). These results suggested that the increased sink strength of LIN, associated with their larger volume, enhanced the overall sink strength within a culm.

Previous studies have indicated that growth duration and dry matter production are potential contributors to stem NSC accumulation (Nagata *et al*., 2002; Takai *et al*., 2006). However, in the current experiments, culm NSC accumulation was not significantly correlated with growth duration, cumulative solar radiation, or aboveground dry matter at the whole-plant level before heading (Supplementary Fig. S10A). Instead, dry matter allocation to the culm in the aboveground organs was a critical factor in determining culm NSC accumulation. The length and volume of LIN were positively correlated with the dry matter ratio allocated to the culm (Supplementary Fig. S10B), indicating that culms with larger LIN volumes received a greater proportion of assimilates. This preferential allocation of assimilates to culms with larger LIN likely explains why the enlargement of LIN promotes NSC accumulation efficiency in UIN, as discussed above. Moreover, such enhanced sink demand by the culm may also mitigate the sugar-mediated feedback inhibition of leaf photosynthesis by facilitating continuous assimilate export from the source leaves, which may lead to increased assimilate allocation to the culm, as proposed in source-sink regulation models (Slewinski, 2012).

Taken together, the cultivars with smaller upper internodes and larger lower internodes exhibited a higher capacity for NSC accumulation in the culm, which could be attributed to the contrasting roles of the upper and lower internodes in regulating NSC storage (Fig. 7).

**Fig. 7.**
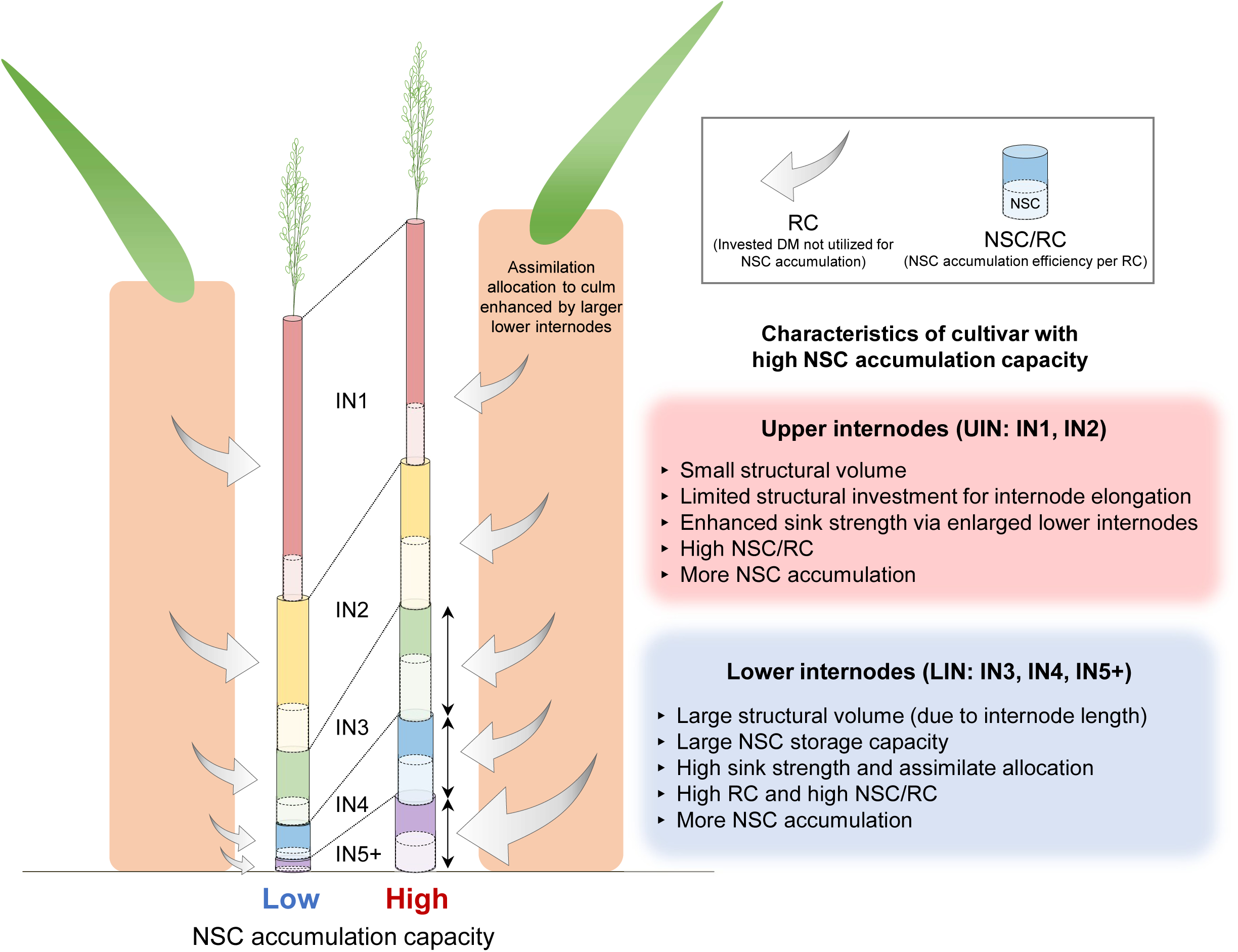
Conceptual model illustrating cultivar differences in morphological traits determining culm NSC accumulation. This diagram summarizes how morphological and physiological characteristics of the upper internodes (UIN; IN1–IN2) and lower internodes (LIN; IN3–IN5+) differentially contribute to culm NSC storage. Cultivars with high NSC accumulation capacity are characterized by smaller UIN volumes and larger LIN volumes. Reduced structural investment in the UIN limits carbon allocation to internode elongation and structural growth, resulting in higher NSC accumulation efficiency (NSC/RC) in these internodes. In contrast, the enlarged LIN provide a large storage capacity for NSC and exhibit high sink strength. The larger volume and enhanced sink strength of the LIN promote greater assimilate allocation to the culm as a whole, thereby increasing sink strength and NSC accumulation efficiency in the UIN.

### VCR as a robust indicator of culm NSC accumulation capacity

To identify a robust morphological indicator associated with NSC accumulation capacity and provide implications for breeding programs, several morphological traits were evaluated (Fig. 4A). The main stem VCR of the UIN/LIN emerged as the most suitable indicator because it consistently showed the strongest correlation with NSC accumulation, not only in the main culm, but also at the whole-plant level (Fig. 4B). This is likely because VCR integrates both the negative contributions of UIN and the positive contributions of LIN, whereas simple volume metrics capture only the total size without reflecting these contrasting functional effects. In addition to its strong association with NSC accumulation, the VCR exhibited high year-to-year reproducibility across experiments (Fig. 4B). This stability indicates that VCR serves as a reliable predictor for evaluating the genetic basis of NSC storage capacity, as it is less affected by environmental fluctuations than NSC accumulation itself (Nagata *et al*., 2001; Wakabayashi *et al*., 2022). Although VCR measurement requires destructive sampling, it can be quantified far more rapidly and with a substantially higher throughput than biochemical NSC analysis, which involves labor-intensive extraction and colorimetric procedures. Accordingly, these advantages indicate that VCR is a practical and robust phenotypic indicator for breeding programs aimed at enhancing stem NSC storage capacity.

### Relationships between leaf sheath NSC accumulation and morphological traits of internodes

Although the NSC content and buffering effect against yield loss were lower in the leaf sheath than in the culm (Supplementary Fig. S6B, S7B), leaf sheaths represent an important temporal NSC storage organ (Nagata et al., 2001; Chen and Wang, 2008). Leaf sheath NSC content showed a strong positive correlation with that of the culm (Supplementary Fig. S6B), suggesting that internode morphological traits also influenced NSC accumulation in leaf sheaths. Active elongation of the UIN was associated with a reduction in NSC content in the leaf sheaths (Supplementary Fig. S9), which was accompanied by changes in the sink strength of the lower leaf sheaths (Fig. 6D). These relationships were also observed among the cultivars, except for one (Supplementary Fig. S11). Leaf sheaths, particularly those at lower positions, transition from sink to source organs earlier than the culm, even at the pre-heading stage (Wakabayashi *et al*., 2022). Therefore, NSC accumulation in these tissues may be particularly sensitive to the assimilation demands generated by the elongation of the upper internodes. In contrast, in the plant growth regulator experiment, the elongation-promoting LIN treatment increased NSC accumulation in the upper leaf sheaths. However, this effect was likely indirect and attributable to the increased length of the upper leaf sheaths (Supplementary Table S7) because the developmental stage of the upper leaf sheaths coincided with the elongation stage of the lower internodes (Wakabayashi *et al*., 2022). In addition, no significant relationship between upper leaf sheath length and LIN length was observed among the cultivars, supporting this interpretation (Supplementary Fig. S12). In contrast, at the cultivar level, genotypes with longer LIN consistently exhibited longer lower leaf sheaths, suggesting that such morphological coordination may influence NSC accumulation in the leaf sheaths (Supplementary Fig. S12, Table S8).

### Agronomic consequences of modifying internode VCR

Culm and internode length are important traits associated with grain yield and biomass production. Therefore, it was assumed that the alteration of internode VCR within a culm also influenced agronomic traits other than NSC accumulation. Several previous studies have indicated that the length of lower internodes negatively affects lodging-related traits in rice (Wu *et al*., 2017; W. J. Zhang *et al*., 2016). However, these studies have primarily evaluated the effects of changes in lower internode length induced by nitrogen fertilization or shading treatments on lodging characteristics. In contrast, studies examining the relationships between lodging-related traits and various morphological traits using multiple cultivars have shown that the length of lower internodes does not affect lodging characteristics (Islam *et al*., 2007; Liu *et al*., 2018). Thus, the negative effects of lower internode elongation on lodging resistance appear to arise primarily from environmentally-induced variation within a cultivar, rather than inherent genotypic differences. Meanwhile, in the present study, IN5+ length was strongly and positively correlated with total culm length (Supplementary Fig. S13), indicating that elongation of the lower internodes accompanied overall culm elongation. Although rice breeding has traditionally favored short-statured cultivars as a primary strategy to improve lodging resistance (Hedden, 2003; Shah *et al*., 2019), the importance of long-culm traits in forage, bioethanol, and high-yield rice cultivars has recently been highlighted (Wu et al., 2011; Okuno et al., 2014; Nomura et al., 2019). Lodging resistance can be achieved even in long-culm cultivars by increasing the culm cross-sectional area and thickness of the cortical fiber tissues (Ookawa *et al*., 2010, 2014). Therefore, when modifying the length of the lower internodes, it is essential to simultaneously enhance culm stiffness to maintain adequate lodging resistance.

In addition to the effects of lower internode or culm length on lodging resistance, panicle size was also associated with internode morphological traits. In particular, the length or diameter of the upper internodes is positively correlated with the spikelet number or panicle length (Yamamoto *et al*., 2001; Sunohara *et al*., 2003; Liu *et al*., 2008; Harshitha *et al*., 2024). In the present study, the diameters of IN1, IN2, and IN3 were positively correlated with spikelet number (Supplementary Fig. S14). Therefore, reducing the volume of the upper internodes through selection for smaller culm diameters may lead to yield penalties due to a reduction in spikelet number. In contrast, no significant correlation was observed between the lengths of IN1 and IN2 and spikelet number. These results suggest that modification of UIN length is an effective approach in enhancing the NSC accumulation efficiency of the upper internodes without negatively affecting panicle size (Sunohara *et al*., 2006).

Variations in internode VCR inevitably alter the vertical distribution of leaves and the overall canopy structure, which may in turn modify the within-canopy light profiles and micrometeorological conditions. Therefore, further analyses that integrate these components are necessary to elucidate how VCR modifications influence yield formation.

In the current study, the volume composition ratio of internodes within a culm was shown to be a key determinant of NSC accumulation capacity in rice. By integrating the contrasting contributions of the upper and lower internodes, VCR provides a robust morphological indicator that is reproducible across environments and cultivars. The findings highlight the importance of internode architecture in regulating carbon allocation within the culm, and offer a practical framework for improving stem carbohydrate storage capacity in rice breeding programs.

## Acknowledgements

We thank the staff at the Institute for Sustainable Agro-ecosystem Services (ISAS), The University of Tokyo, for technical support in rice cultivation and management.

## Author contributions

Y.W. designed and conducted the experiments, analyzed the data, and prepared the manuscript. Y.W. and M.S. performed the quantification of soluble sugars and starch and measured AGP activity. All authors read and approved the final manuscript.

## Conflict of interesth

The authors declare that they have no competing interests.

## Data availability

All data supporting the findings of this study are available from the corresponding author upon reasonable request.

## Abbreviations

LIN: lower internodes
NSC: non-structural carbohydrates
RC: residual components
UIN: upper internodes
VCR: volume composition ratio

## Supplementary figure legends

Supplementary Fig. S1. Experimental design of plant growth regulator treatments targeting specific internodes.

(A) Summarized treatment combinations. (B) The diagram illustrates the developmental progression of internodes (IN1–IN5) and corresponding leaf sheaths (LS1–LS5) from panicle formation to heading. Plant growth regulator treatments were applied at 25 days before heading (DBH) and 3 DBH to modify elongation of IN4 (representing the lower internodes) and IN2, respectively.

Supplementary Fig. S2. Workflow for calculating internode cross-sectional traits from scanned images. The procedure used to obtain internode diameter, wall thickness, and cross-sectional area from scanned cross-sectional images is illustrated. (1) Image thresholding: The raw cross-sectional image is binarized to detect the tissue boundaries in the outer or inner circles. (2) Area measurement: The areas of the outer (red) and inner (green) regions are measured. The cross-sectional area is calculated as the difference between these two areas. (3) Reconstruction of concentric annuli: Idealized concentric circles equivalent in area to the measured outer and inner regions are reconstructed to minimize irregularities caused by tissue deformation or sectioning artifacts. (4) Calculation of structural traits: From the reconstructed circles, the diameter (outer circle) and wall thickness (difference between outer and inner radii) are calculated.

Supplementary Fig. S3. Panicle number per plant for each cultivar evaluated in the 2022 field experiment. Bars show means ± SE (*n* = 4). Different letters above the bars indicate significant differences among cultivars (Tukey’s HSD test, *P* < 0.05).

Supplementary Fig. S4. Solar radiation and mean air temperature during the pre-heading period for each cultivar grown in 2022 and 2024.

(A, B) Daily solar radiation (MJ m□² d□¹) averaged over the period from panicle formation to heading in 2022 (A) and from panicle formation to full heading in 2024 (B). (C, D) Mean air temperature (°C) during the same period in 2022 (C) and 2024 (D). Bars show means ± SE (*n* = 4). Different letters above the bars indicate significant differences among cultivars (Tukey’s HSD, *P* < 0.05). *F* values from one-way ANOVA are shown above each panel.

Supplementary Fig. S5. Relationships between culm NSC content and climatic conditions before heading. (A) Relationship between solar radiation and main culm NSC content in 2022. (B) Relationship between solar radiation and whole-plant culm NSC content in 2024. (C) Relationship between mean air temperature and main culm NSC content in 2022. (D) Relationship between mean air temperature and whole-plant culm NSC content in 2024. Regression lines are shown with 95% confidence intervals (shaded). Correlation coefficients (*r*) and *P*-values are from Pearson’s tests (*n* = 10).

Supplementary Fig. S6. Leaf sheath NSC content and its association with culm NSC content in rice cultivars.

(A) Leaf sheath NSC content of each cultivar in 2022 (left) and 2024 (right). In 2022, leaf sheath NSC content is shown as stacked bars separating LS1–LS2 and LS3+ (leaf sheaths at and below LS3), whereas in 2024 total leaf sheath NSC content is shown. Bars show means ± SE (n=4). Different letters above bars indicate significant differences among cultivars (Tukey’s HSD, *P* < 0.05). (B) Relationships between leaf sheath NSC content and culm NSC content in 2022 (left) and 2024 (right). Black solid lines and gray shaded areas indicate the regression lines and 95% confidence intervals calculated using all cultivars, whereas red solid lines and red shaded areas indicate those calculated using eight cultivars excluding MR and HA. Pearson’s correlation coefficients (*r*), *p-*values, coefficients of determination (*R²*), and regression equations are shown in each panel.

Supplementary Fig. S7. Relationship between culm or leaf sheath NSC and yield reduction under defoliation.

(A) Reduction ratio of grain dry weight (DW) under defoliation treatment of each cultivar. Bars show means ± SE (*n*=4). Different letters above bars indicate significant differences among cultivars (Tukey’s HSD, *P* < 0.05). The reduction ratio was calculated as grain DW under defoliation relative to the non-defoliated control. (B) Relationships between the reduction ratio of grain DW under defoliation and NSC content at the whole-plant level in the culm (left) and the leaf sheath (right). Regression lines are shown with 95% confidence intervals (shaded). Correlation coefficients (*r*) and *P*-values are from Pearson’s tests (*n* = 10).

Supplementary Fig. S8. Relationships between the upper internodes (UIN) VCR and culm NSC accumulation, and year-to-year reproducibility.

Relationships between UIN VCR and main culm NSC content in 2022 (left), between UIN VCR and whole-plant culm NSC content in 2024 (middle) and between UIN VCR in 2022 and 2024 (right), showing year-to-year stability. Regression lines are shown with 95% confidence intervals (shaded). Correlation coefficients (*r*) and *P*-values are from Pearson’s tests (*n* = 10).

Supplementary Fig. S9. Effects of internode morphological modification on leaf sheath NSC accumulation.

Leaf sheath (LS) NSC content in the whole leaf sheath (LS) (top), upper leaf sheaths (LS1, LS2) (middle), and lower leaf sheaths (LS3–LS6) (bottom) under different plant growth regulator treatments. Bars show means ± SE (n = 5). Horizontal brackets indicate statistically significant differences among treatments (Dunnett vs. MM, * P < 0.05, ** P < 0.01).

Supplementary Fig. S10. Relationships between culm NSC and traits associated with growth duration or dry matter production/allocation.

(A) Correlation pathways showing the relationships of culm NSC content with growth duration, cumulative solar radiation (SR), aboveground dry matter (DM), and culm DM ratio before heading (in 2022: left) or full-heading (in 2024: right). Red line indicates positive correlations. Line thickness corresponds to correlation strength. Significance levels: ** *P* < 0.01, *** *P* < 0.001 (*n* = 10). (B) Relationships between culm DM ratio and the length (top) or volume (bottom) of LIN in 2022 (left) and 2024 (right). Regression lines are shown with 95% confidence intervals (shaded). Correlation coefficients (*r*) and *P*-values are from Pearson’s tests (*n* = 10).

Supplementary Fig. S11. Relationships between length or volume of upper internodes (UIN) and leaf sheath NSC content.

Relationships between length (top) or volume (bottom) of UIN and NSC content of the whole leaf sheaths (LS) (left), the upper leaf sheaths (LS1, LS2) (middle) and the lower leaf sheaths (LS3+) (right). Black lines and gray shaded areas indicate the regression lines and 95% confidence intervals calculated using all cultivars, whereas blue lines and shaded areas indicate those calculated using nine cultivars excluding MR. Pearson’s correlation coefficients (*r*), *P*-values, coefficients of determination (*R²*), and regression equations are shown in each panel.

Supplementary Fig. S12. Correlation matrix of leaf sheath NSC, leaf sheath length, and internode morphology in 2022.

Correlation matrix among NSC content in leaf sheaths, leaf sheath length, and morphological traits of the upper and lower internodes in 2022. Leaf sheath NSC content was evaluated for all leaf sheaths combined (LS), LS1-2, and LS3 and below (LS3+). Leaf sheath length was analyzed for individual leaf sheaths (LS1–LS4), and LS5+ represents the mean length of leaf sheaths at and below LS5. Internode morphological traits were summarized as length and volume of the upper internodes (UIN; IN1–IN2) and lower internodes (LIN; IN3–IN5+). Correlation analyses were performed using data from nine cultivars, excluding MR. Circle size and color indicate correlation strength and direction, respectively (Pearson’s *r*). Significance levels: † *P* < 0.10, * *P* < 0.05, ** *P* < 0.01, *** *P* < 0.001 (*n* = 9).

Supplementary Fig. S13. Relationship between lower internode length (LIN) and culm length. Relationships between LIN length and culm length of main stem in 2022 (left) and in 2024 (right). Regression lines are shown with 95% confidence intervals (shaded). Correlation coefficients (*r*) and *P*-values are from Pearson’s tests (*n* = 10).

Supplementary Fig. S14. Yield component traits and their relationships with internode morphology in 2024. (A) Spikelet number per plant (left), spikelet number per panicle (middle), and panicle number per plant (right) of each cultivar in 2024. Bars show means ± SE (n=4). Different letters above bars indicate significant differences among cultivars (Tukey’s HSD, P < 0.05). (B) Correlation matrix between yield components and internode morphological traits (VCR, volume, length, cross-sectional area, diameter, and wall thickness). Circle size and color indicate correlation strength and direction, respectively (Pearson’s r). Significance levels: † P < 0.10, * P < 0.05 (n = 10).

## Supplementary table legends

Supplementary Table. S1. Cultivar information and phenological traits in the 2022 and 2024 field experiments.

Cultivars were classified into indica (IND), temperate japonica (TEJ), or japonica–indica admixture (MIX) based on population structure analysis reported by Yonemaru *et al*. (2014). Values indicate means (*n* = 4). Different letters within each column indicate significant differences among cultivars (Tukey’s HSD, *P* < 0.05). *F* values and significant levels are from one-way ANOVA. Significance levels: *** *P* < 0.001.

Supplementary Table. S2. NSC-related traits across cultivars in 2022 and 2024 field experiments.

Values indicate means (*n* = 4). Different letters within each column indicate significant differences among cultivars (Tukey’s HSD, *P* < 0.05). *F* values and significant levels are from one-way ANOVA. Significance levels: ** *P* < 0.01, *** *P* < 0.001.

Supplementary Table. S3. Morphological traits of individual internodes and culm across cultivars in the 2022 field experiment.

Values indicate means (*n* = 4). Different letters within each column indicate significant differences among cultivars (Tukey’s HSD, *P* < 0.05). *F* values and significant levels are from one-way ANOVA. Significance levels: ** *P* < 0.01, *** *P* < 0.001; n.s. not significant.

Supplementary Table. S4. Morphological traits of individual internodes and culm across cultivars in the 2024 field experiment.

Values indicate means (*n* = 4). Different letters within each column indicate significant differences among cultivars (Tukey’s HSD, *P* < 0.05). *F* values and significant levels are from one-way ANOVA. Significance levels: ** *P* < 0.01, *** *P* < 0.001.

Supplementary Table. S5. NSC-related traits of individual internodes and culm across treatments in the plant growth regulator experiment.

Values indicate means (*n* = 5). Symbols indicate significant differences from the control (MM) (Dunnett’s test). *F* values and significant levels are from one-way ANOVA. Significance levels: † *P* < 0.10, * *P* < 0.05, ** *P* < 0.01, *** *P* < 0.001; n.s. not significant.

Supplementary Table. S6. Morphological traits of individual internodes and culm across treatment in the plant growth regulator experiment.

Values indicate means (*n* = 5). Symbols indicate significant differences from the control (MM) (Dunnett’s test). *F* values and significant levels are from one-way ANOVA. Significance levels: † *P* < 0.10, * *P* < 0.05, ** *P* < 0.01, *** *P* < 0.001; n.s. not significant.

Supplementary Table. S7. Length of individual leaf sheaths across treatments in the plant growth regulator experiment.

Values indicate means (*n* = 5). Symbols indicate significant differences from the control (MM) (Dunnett’s test). *F* values and significant levels are from one-way ANOVA. Significance levels: * *P* < 0.05, ** *P* < 0.01, *** *P* < 0.001; n.s. not significant.

Supplementary Table. S8. Length of individual leaf sheaths across cultivars in the 2022 field experiment. Values indicate means (*n* = 4). Different letters within each column indicate significant differences among cultivars (Tukey’s HSD, *P* < 0.05). *F* values and significant levels are from one-way ANOVA. Significance levels: *** *P* < 0.001; n.s. not significant.

